# A role for JAK2 in mediating cell surface GHR-PRLR interaction

**DOI:** 10.1101/2023.09.01.555812

**Authors:** Chen Chen, Jing Jiang, Tejeshwar C. Rao, Ying Liu, Tatiana T. Marquez Lago, Stuart J. Frank, André Leier

## Abstract

Growth hormone (GH) receptor (GHR) and (full-length) prolactin (PRL) receptor (PRLR) are transmembrane class I cytokine receptors that co-exist in various normal and cancerous cells. Both receptors respond to their associated ligands predominantly by activating the Janus Kinase 2 (JAK2)-signal transducer and activator of transcription (STAT) signaling pathways, and both are also known to initiate receptor-specific JAK2-independent signaling. Together with their cognate ligands, these receptors have been associated with pro-tumorigenic effects in various cancers, including breast cancer (BC). Human GH is known to bind GHR and PRLR, while PRL can only bind PRLR. A growing body of work suggests that GHR and PRLR can form heteromers in BC cells, modulating GH signal transduction. However, the dynamics of PRLR and GHR on the plasma membrane and how these could affect their respective signaling still need to be understood.

To this end, we set out to unravel the spatiotemporal dynamics of GHR and PRLR on the surface of human T47D breast cancer cells and γ2A-JAK2 cells. We applied direct stochastic optical reconstruction microscopy (dSTORM) and quantified the colocalization and availability of both receptors on the plasma membrane at the nanometer scale at different time points following treatment with GH and PRL. In cells co-expressing GHR and PRLR, we surprisingly observed that not only GH but also PRL treatment induces a significant loss of surface GHR. In cells lacking PRLR or expressing a mutant PRLR deficient in JAK2 binding, we observed that GH induces downregulation of cell surface membrane-bound GHR, but PRL no longer induces loss of surface GHR. Colocalizations of GHR and PRLR were confirmed by proximity ligation (PL) assay.

Our results suggest that PRLR-GHR interaction, direct or indirect, is indispensable for PRL- but not GH-induced loss of surface GHR and for both GH-induced and PRL-induced increase of surface PRLR, with potential consequences for downstream signaling. Furthermore, our results suggest that JAK2 binding via the receptor intracellular domain’s Box1 element is crucial for the observed regulation of one class I cytokine receptor’s cell surface availability via ligand-induced activation of another class I cytokine receptor. Our findings shed new light on the reciprocal and collective role that PRLR and GHR play in regulating cell signaling.

## Introduction

Growth hormone (GH) and prolactin (PRL) are hormones emanating mainly from the anterior pituitary. The primary function of GH is regulating anabolism and metabolism [1, 2], while PRL has important roles in breast development and lactation [3]. There is mounting evidence pointing at both hormones and their receptors playing roles in various types of cancer [4–11], including breast cancer (BC) [12–17], where GHR is frequently present and PRLR is often found overexpressed [18–25]. While they have been mostly associated with pro-tumorigenic effects, PRL has also been reported to show anti-tumor effects and, like PRLR, has been associated with good prognosis in certain BC subtypes [26–29]. However, a humanized neutralizing monoclonal antibody directed against the extracellular domain of PRLR showed no anti-tumor effect when administered in patients with PRLR-positive metastatic BC [30]. This suggests that PRLRs’ pro-tumorigenic function may not be as relevant as previously thought or depends on other circumstances such as the presence or absence of other hormone receptors, with which they may interact.

Both GH receptor (GHR) and PRL receptor (PRLR) are structurally similar transmembrane glycoproteins and belong to the class I cytokine receptor superfamily [31, 32]. As is characteristic of the majority of their superfamily members, both receptors contain a proline-rich Box1 motif in the membrane-proximal region of their intracellular domains (ICDs), which is followed by a short inter-box region, and a less conserved Box2 motif that is rich in acidic and aromatic residues [33–35]. For both receptors different isoforms have been identified: GHR has three additional isoforms two of which are created through alternative splicing events and lack the Box1-Box2 motif [36] and the third is the exon 3 deleted GHR (d3-GHR). Aside from the PRLR long (or full length) isoform (LF), other membrane-bound isoforms have been identified, including a long form lacking domain D1 (ΔS1), an intermediate form (IF), and the short forms S1a, S1b, and S1c [3, 37]. All isoforms except for S1b and S1c contain the Box1-Box2 motif, but S1b and S1c still contain the Box1 motif. If not specifically stated otherwise, we will refer here by default to the receptors’ full-length (or long) isoforms.

GH can bind and introduce a conformational change to both GHR and PRLR, allowing receptor activation and downstream signaling [38–42], but, unlike GH, PRL can only bind to PRLR [41, 43–45]. Both GHR and PRLR lack intrinsic kinase activity. Following ligand binding, downstream signal transduction involves predominantly activating the associated cytoplasmic tyrosine kinase, Janus kinase 2 (JAK2), bound to the receptors’ Box1 elements. This is followed by phosphorylation of the signal transducer and activator of transcription 5 (STAT5a and STAT5b) [46, 47], although other receptor-specific JAK2-independent signal transduction pathways may also be activated. Of note, JAK2 is not only activating the canonical JAK2- STAT5 pathway, but it can also signal via STAT1 and STAT3 [48–51] and activate non-canonical kinase pathways such as Ras/Raf/MAPK/ERK [52–54] and PI3K/Akt/mTOR [55–58], as well as influence gene expression through histone modification [59, 60]. Moreover, activation of PRLR downstream pathways is isoform-specific, with the short isoforms activating the MAPK and PI3K pathways [61–63]. For comprehensive reviews of JAK2-independent signaling pathways, we refer the reader to [64, 65].

Increasing evidence indicates that GHR and PRLR interact. Two decades ago, it was shown that ovine GHR (oGHR) and PRLR (oPRLR) can tightly associate with each other following stimulation with placental lactogen [66]. These studies utilized chimeric receptors consisting of the extracellular domain (ECD) of human granulocyte and macrophage colony-stimulating factor (hGM-CSF) receptor (hGM-CSFR) along with a part of either oGHR ICD or oPRLR ICD. After hGM-CSF treatment of cells co-expressing oGHR chimera and oPRLR chimera, JAK2 was effectively activated, and protein-protein interaction of both chimeric receptors was detected via co-immunoprecipitation [66, 67]. Additionally, our previous work revealed a specific ligand-independent human GHR (hGHR) - human PRLR (hPRLR) association in human T47D breast cancer cells, which endogenously express both receptors [68]. Further, the use of split luciferase complementation assays has suggested that hGHR homodimers and hPRLR homodimers form hGHR/hPRLR multimers [69] and extracellular subdomain 2 of the hGHR or hPRLR determines the dimerization partner [70]. Although biochemical studies and luciferase complementation assays strongly support the notion that an interplay between hGHR and hPRLR exists, the observed outcomes are ascribed to total receptors within cells, irrespective of subcellular localization.

In the present study, we directly visualized the cell surface interaction of hGHR and hPRLR and how it changes upon ligand treatment. Specifically, we used direct stochastic optical reconstruction microscopy (dSTORM) [71] to visualize single receptor clusters of hGHR and hPRLR on cell surfaces. In dSTORM, a super-resolution microscopy technique, individual fluorophores cycle through reversible transitions between a dark and a fluorescent state [72]. Thus, a fluorophore emits photons multiple times before permanently being photobleached. These blinking events and their localizations are recorded and, although the exact number of proteins in clusters is difficult to determine, the number of localizations is strongly correlated with the number of receptors [73, 74]. Thus, the dSTORM approach can deliver high-resolution images to reveal the localization or arrangement of individual membrane receptor systems providing valuable insight into their interactions with other proteins at the cell surface. Given that receptors are highly trafficked to and from the cell surface and to avoid signal detection from cytosolic receptors, we used monoclonal antibodies to distinctly label the extracellular S2 domain of GHR and PRLR on non-permeabilized cells. Descriptive spatial analysis using Ripley’s K- and L-function [75] indicates that both hGHR and hPRLR are organized in nanometer-scale clusters on the T47D cell surface. To further gain quantitative information about GHR and PRLR nanoclusters, we applied DBSCAN (density-based spatial clustering of applications with noise) [76]. Subsequently, individual cluster contours were delineated, individual clusters were assigned a cluster ID, and receptor abundance was analyzed separately for each cluster and receptor. By doing so, we detected and calculated homomeric and heteromeric hGHR and hPRLR clusters on cell surfaces. Colocalizations of GHR and PRLR were also confirmed by proximity ligation (PL) assay. Lastly, we explored which receptor domains determine the interaction of hGHR and hPRLR by creating different truncated or modified hPRLR variants. Our findings indicate that the intracellular Box1 region is an essential determinant of hGHR and hPRLR association, suggesting JAK2 may play an important role in the observed ligand-induced and PRLR-mediated downregulation of cell surface GHR – which could also elicit or add to PRLR’s observed anti-tumor effect.

## Results

### Ligands induce an increase in PRLR localizations and a decrease in GHR localizations at the surface of T47D cells

To investigate the dynamic spatial distribution of endogenous hGHR and hPRLR on the surface of breast cancer cells, we used dSTORM under the total internal reflection fluorescence (TIRF) illumination mode [77]. The dSTORM images show that hGHR and hPRLR form nanometer-scale clusters and are broadly distributed on the surface of T47D cells (Fig. 1). To assess the ligands’ effects on the spatial distribution of hGHR and hPRLR on the cell surface, we conducted time-course experiments using T47D cells with human GH or human PRL (500 ng/ml each). Such stimulation is known to induce rapid and substantial STAT5 phosphorylation [68]. We first analyzed the cell surface localization density (number of localizations per μm^2^) of hGHR or hPRLR in both resting and ligand-stimulated conditions. The abundance of surface-hPRLR was rapidly increased, reaching its maximum after 3 min of GH or PRL treatment with a ∼5.6-fold and ∼4.5- fold increase compared to the basal value, respectively (Fig. 2A and 2B). This increase was followed by a rapid decline: after GH treatment for 5 min, the localization density dropped from 200.6 ± 14.7 per μm^2^ to 57.2 ± 3.6 per μm^2^; after PRL treatment for 5 min, the density fell from 159.4 ± 16.2 per μm^2^ to 94.8 ± 16.0 per μm^2^. In contrast to hPRLR, the density of hGHR significantly decreased from 43.3 ± 5.5 per μm^2^ basally to 25.8 ± 2.7 per μm^2^ after 3min of GH stimulation. After 5 min and 10 min of GH stimulation, the density of hGHR remained relatively low at 26.6 ± 2.3 per μm^2^ and 19.3 ± 1.5 per μm^2^, respectively (Fig. 2C). Surprisingly, PRL, which does not bind to hGHR, also induced a loss of surface hGHR on T47D cells. After 1 min of PRL treatment, only 36% of hGHR remained on the cell surface compared to the basal state, and hGHR density remained low for at least 10 min (Fig. 2D). A schematic illustration of GH- and PRL-induced hGHR and hPRLR density changes is shown in Fig. 2E. We previously observed that hGHR and hPRLR specifically co-immunoprecipitate in the absence of added ligands in T47D cells [68]; thus, we postulate that the propensity of hGHR and hPRLR to physically interact underlies our observed loss of hGHR in response to PRL stimulation.

**Fig. 1.**
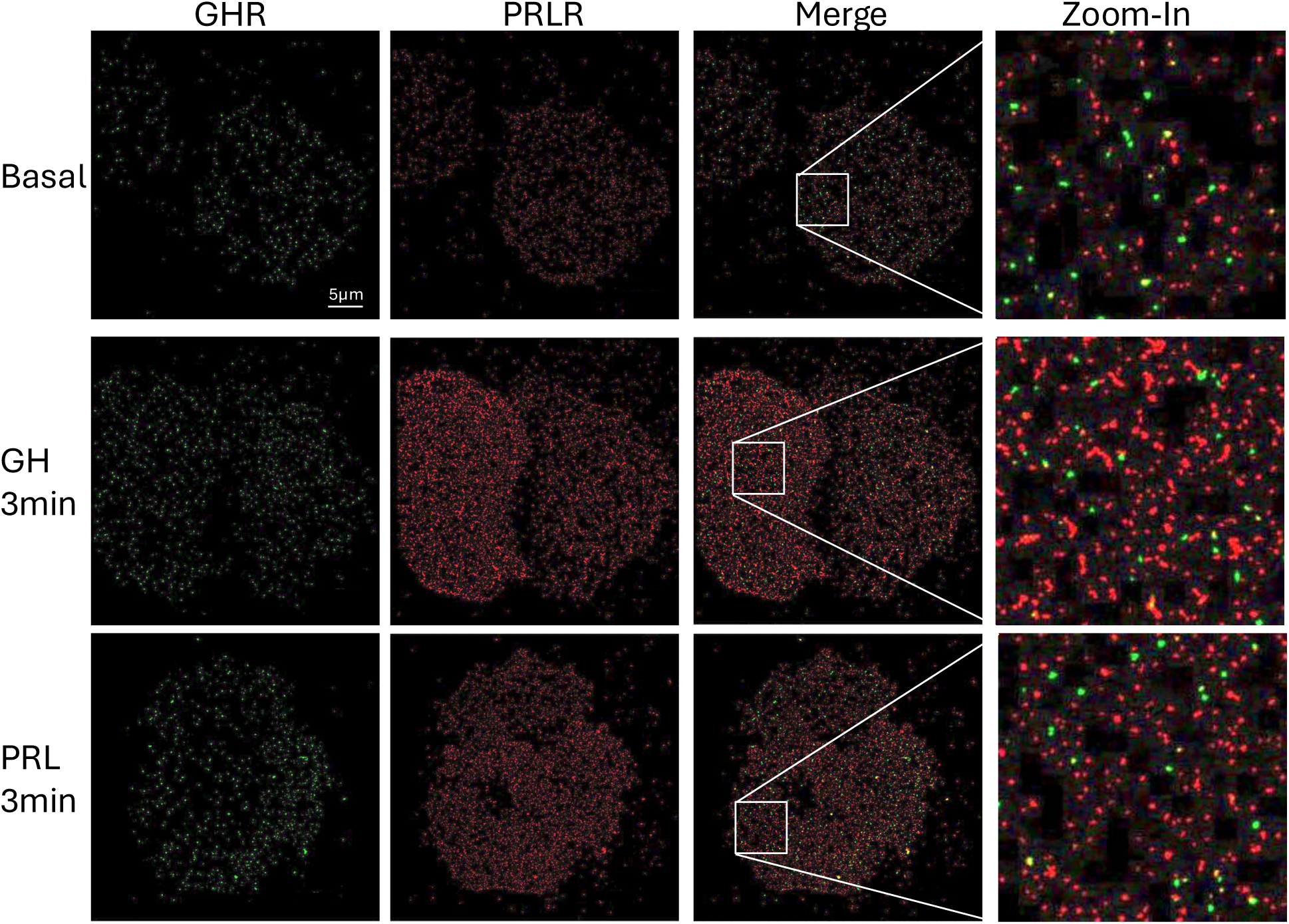
Ligand-induced increase of PRLR localizations and decrease of GHR localizations at the surface of T47D cells. A representative set of reconstructed dSTORM images, showing Gaussian-fitted localizations obtained from fluorophore blinking events. GHR is labeled with Alexa 568-conjugated antibody (green, channel 1), and PRLR is labeled with Alexa 647-conjugated antibody (red, channel 2). The last column of images shows the merging of these two channels. T47D cells were either left untreated (upper row), exposed to 500ng/mL GH for 3 min (middle row), or exposed to 500ng/mL PRL for 3 min (lower row). Brightness was increased by 40% and Contrast was reduced by 40% to increase visibility. The images in the first three columns (GHR, PRLR, Merge) have the same scale. The scale bar (5μm) is shown in the upper left image. The images in the right most column are zoomed-in sections from the corresponding merged images.

**Fig. 2.**
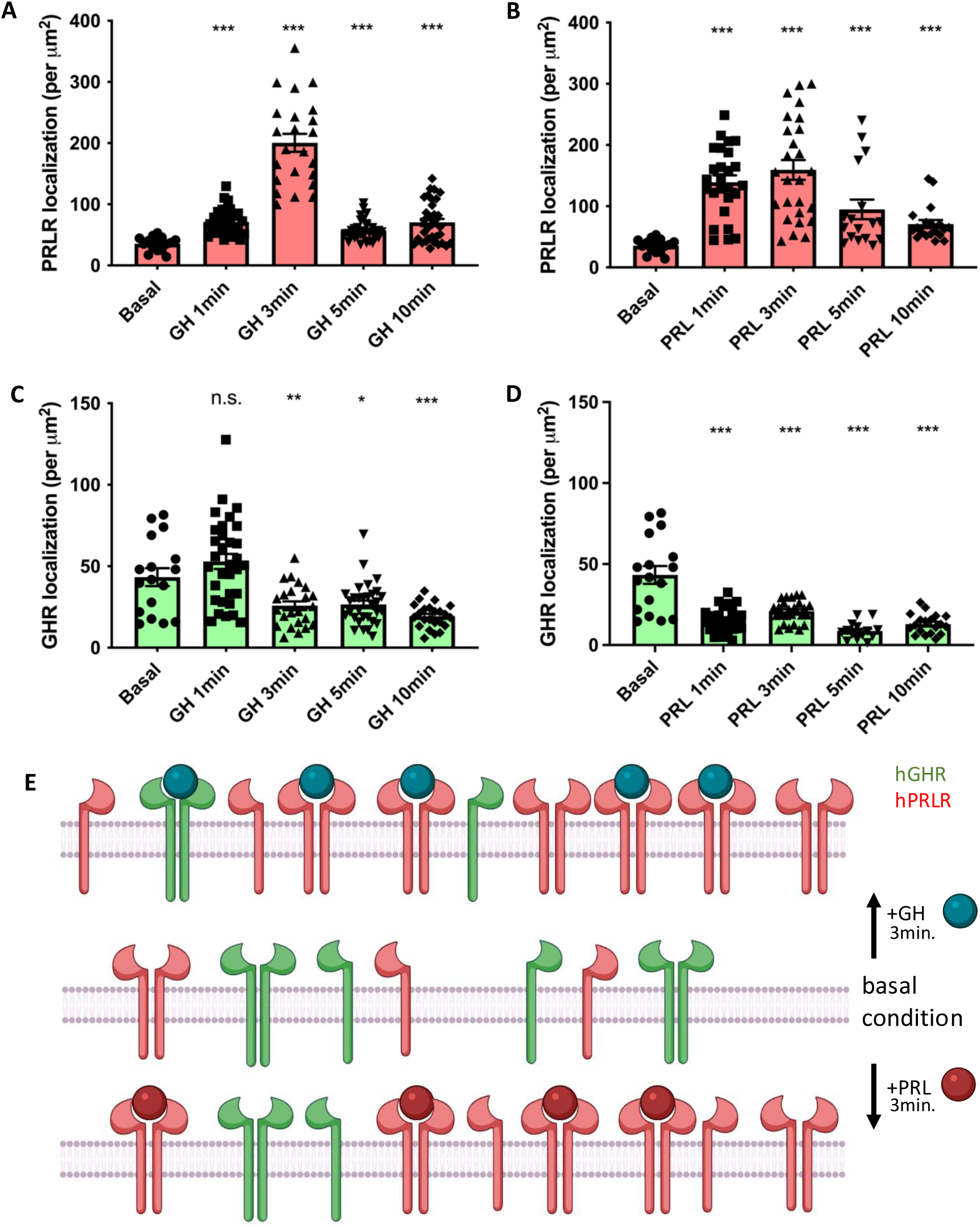
**(A,B)** Quantification of GH-induced (500 ng/ml) (A) and PRL-induced (500 ng/ml) (B) changes of PRLR localizations on the cell surface. **(C, D)** Quantification of GH-induced (C) and PRL-induced (D) changes of GHR localizations on the cell surface. Each data point represents the density of receptors in a 6.25 μm X 6.25 μm area. Data are collected from at least 6 cells from each group and displayed as mean ± SE. * (P<0.05), ** (P<0.01), and *** (P<0.001) indicate the statistical significance in comparison with Basal and are calculated by two-tailed t-tests assuming unequal variances. **(E)** Model depicting hPRLR and hGHR densities and activation schemes in T47D cells. The middle of the three states depicts the basal condition.

### GH or PRL stimulation induces a redistribution of hGHR and hPRLR clusters

We analyzed the dSTORM images using the DBSCAN algorithm to identify different clusters and determine the number of receptor localizations within a cluster (termed ‘cluster size’) (Fig. 3A). We then performed a localization distribution analysis and plotted a histogram of the relative frequency of localizations (termed ‘distribution plot’ in this study). Representative dSTORM images of hGHR as well as associated distribution plots are shown in Fig. 3B.

**Fig. 3.**
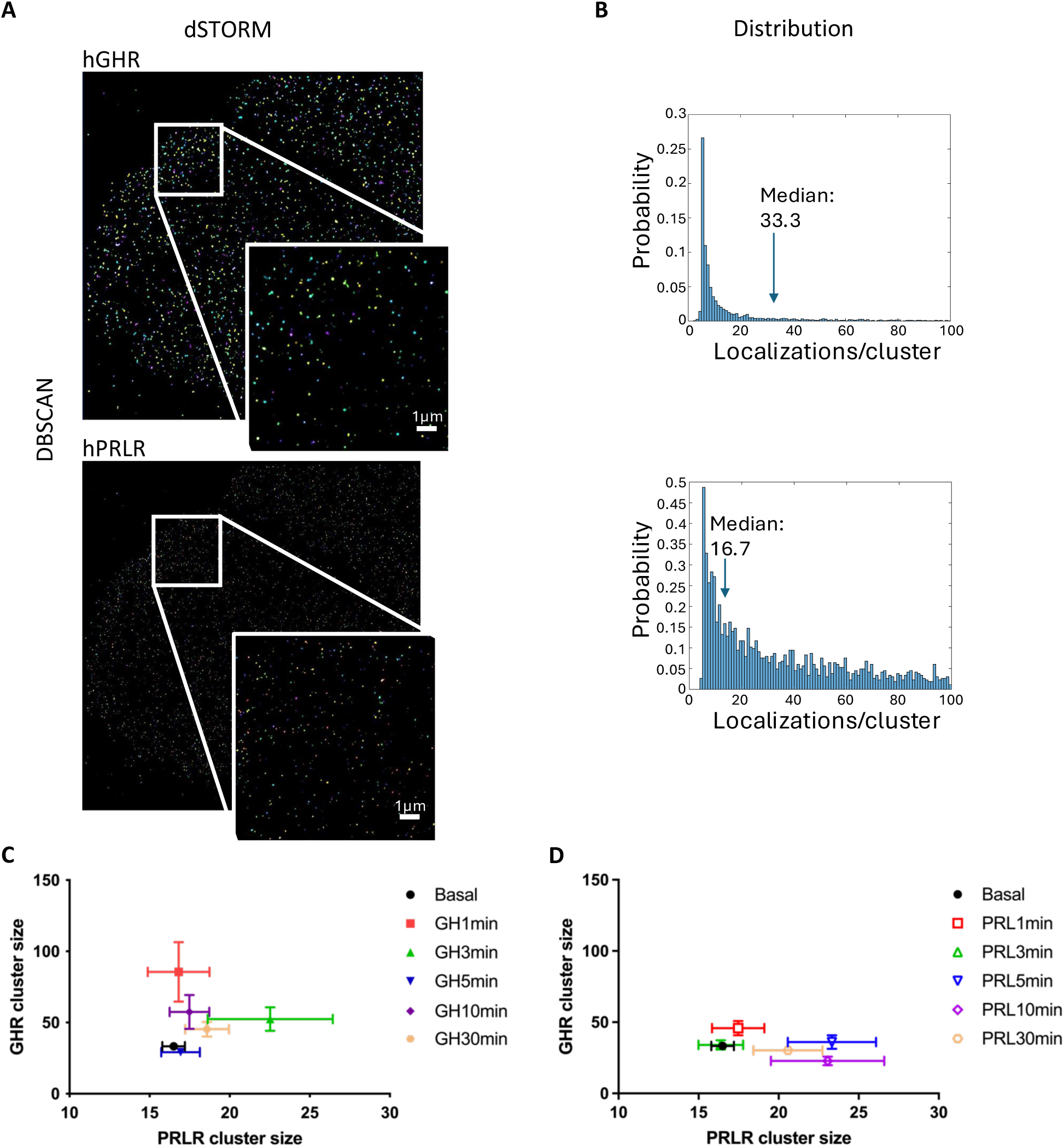
GH/PRL stimulation induces a redistribution of hGHR and hPRLR clusters. **(A)** Sample dSTORM images and application of DBSCAN to identify and measure clusters (scale bar, 1 μm). To distinguish different clusters, each cluster has a different color assigned. **(B)** For similar images corresponding histograms of the relative frequencies of localizations are shown. Median cluster sizes are indicated. **(C, D)** The medians of hGHR and hPRLR cluster sizes following GH (C) and PRL (D) treatment are summarized in these plots. The x-axis and y-axis represent hPRLR and hGHR cluster sizes (counts), respectively. Data are collected from at least 6 cells from each group and displayed as median ± SE.

To gain a better understanding of the changes in GHR-PRLR colocalizations, we obtained and analyzed the corresponding bivariate cluster size distributions. Their median values are summarized in Fig. 3C and 3D. As can be seen, hGHR responds quickly and transiently to GH stimulation by forming larger clusters. After GH treatment for 1 min, the median number of hGHR blinking events in a cluster reaches its peak and amounts to 85.6 ± 20.9 localizations in comparison with 33.3 ± 1.7 localizations per cluster in the basal state (P value = 0.0035). After 5 min, this number is reduced approximately to its pre-stimulation value (29.1 ± 2.2) but appears to continue to fluctuate for at least another 25 minutes (Fig. 3C). In turn, hPRLR responds to GH in a less prominent manner: The median number of hPRLR blinking events in a cluster in the basal state is 16.5 ± 0.7. After 3 min of GH stimulation, the median of PRLR cluster size reaches its maximum, which is 22.5 ± 3.9 (P value = 0.1496), and afterward declines to its basal level (Fig. 3C).

Like the response to GH, hGHR reacts also quickly to PRL treatment. The median number of hGHR blinking events in a cluster culminates at 1 min of PRL treatment with 45.8 ± 5.0 (compared with basal level, P value = 0.0098). In distinction, hPRLR response to PRL is slower. The median number of hPRLR blinking events in a cluster reaches its peak at 5 min of PRL treatment with a median number of 23.3 ± 2.8 (compared with basal level, P value = 0.0233), and after 10 min the median is only slightly less than that maximum (Fig. 3D). Together, these results indicate that upon ligand stimulation hGHR cluster sizes increase transiently and significantly, while changes of hPRLR cluster sizes occur slowly and more subtly.

### Spatial proximity of hGHR and hPRLR upon ligand stimulation

Nanoscale interactions of hGHR and hPRLR on the cell surface have yet to be well established. To evaluate the extent of hGHR and hPRLR surface colocalization on T47D cells, we utilized proximity ligation assays (PLAs). In PLAs, a positive signal appears only when two target proteins are in proximity (<40 nm). Notably, individual treatment with GH or PRL for 5 min decreased the PLA signal observed in untreated cells by 34.4% and 28.1%, respectively (Fig. 4A), suggesting either ligand caused a reduction in the total number of colocalized hGHR and hPRLR clusters. To further validate these observations, we calculated the ratio of colocalized clusters in dSTORM images. Treatment with GH or PRL for 1 min reduced the proportion of colocalized clusters by nearly 50% (Fig. 4B and 4C). To analyze their compositions, we plotted 3D distributions for the colocalized clusters after ligand treatments (Fig. S1A and S1B). The probability of observing co-localized clusters with numbers of hGHR and hPRLR that fall into certain bins is identified by color and by the height of the bar on the z-axis. Following GH treatment for 1 min and 3 min, the number of smallest clusters decreased, and the number of medium-sized clusters increased, suggesting a shift of the bivariate distribution of co-localized cluster sizes toward medium-sized clusters (Fig. S1A). The distribution of colocalized clusters after 5 min of GH treatment is similar to that at the basal state. In contrast, after treatment with PRL for 3 min or more, the number of medium to large-sized clusters, majoritarily containing either GHR or PRLR, decreased, while the number of smallest clusters increased (Fig. S1B). Together, these results demonstrate that hGHR and hPRLR are spatially accessible to each other and form receptor complexes on the T47D cell surface and that the nature of these complexes changes differentially depending on the stimulating ligand.

**Fig. 4.**
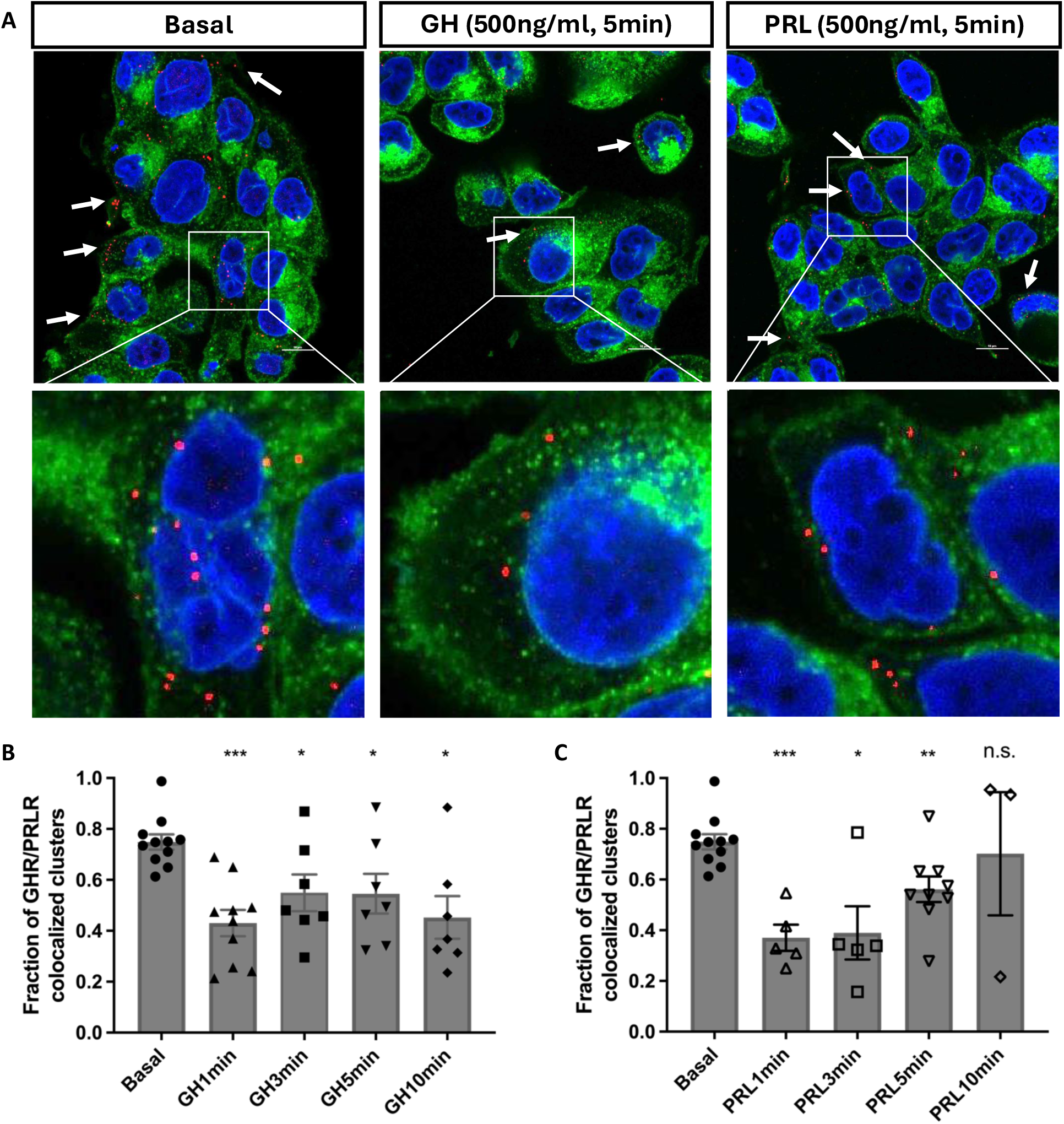
Spatial proximity of hGHR and hPRLR upon ligand stimulation. **(A)** Proximity ligation assays were performed in T47D cells (scale bar, 10 μm). Here we show representative confocal microscopy images of T47D cells at resting and ligand stimulation conditions. The PLA signal (red dots) corresponds to hGHR and hPRLR complexes. Wheat germ agglutinin staining (green) was used to label glycoproteins for cell membrane imaging. DAPI (blue staining) corresponds to cell nuclei. Zoom-in views are provided in the second row. **(B, C)** hGHR and hPRLR colocalization changes in T47D cell surface following 500 ng/ml GH (B) or PRL (C) treatment obtained from dSTORM images using DBSCAN. Each data point represents a ratio of the number of clusters containing both hGHR and hPRLR, over the total number of clusters on the cell surface. Data are displayed as mean ± SE. * (P<0.05), ** (P<0.01), and *** (P<0.001) indicate the statistical significance in comparison with Basal and are calculated by a two-tailed t-test assuming unequal variance.

### Reduction of hGHR induced by PRL on the cell surface is dependent on the presence of PRLR

PRL strongly binds PRLR but not GHR. Yet, PRL induces a decrease of hGHR on the surface of cells expressing both hGHR and hPRLR. Thus, we sought to investigate the effect of PRL on hGHR in the absence of hPRLR. We utilized CRISPR/Cas9 technology to generate hPRLR knockout T47D cells (termed T47D_ΔPRLR_). In addition, to evaluate isolated hPRLR responses to ligands, we also generated hGHR knockout T47D cells (termed T47D_ΔGHR_). Immunoblot analysis with specific GHR and PRLR antibodies confirmed the absence of hPRLR in T47D_ΔPRLR_ cells and hGHR in T47D_ΔGHR_ cells (Fig 5A). Similar to our results with parental T47D cells, GH treatment of T47D_ΔPRLR_ cells rapidly reduces the density of surface hGHR, suggesting that hPRLR need not be present to allow this GH-induced effect (Fig 5B). However, contrary to findings in parental T47D cells, PRL treatment of T47D_ΔPRLR_ cells fails to modulate hGHR surface density (Fig 5C). Like parental T47D cells, treatment of T47D_ΔGHR_ cells with GH (Fig 5D) or PRL (Fig 5E) yielded increased hPRLR surface localizations. Thus, we conclude that the PRL-induced decrease of hGHR in T47D cells is dependent on the presence of hPRLR, but the ability of both GH and PRL to increase surface hPRLR in T47D cells is independent of hGHR’s presence.

**Fig. 5.**
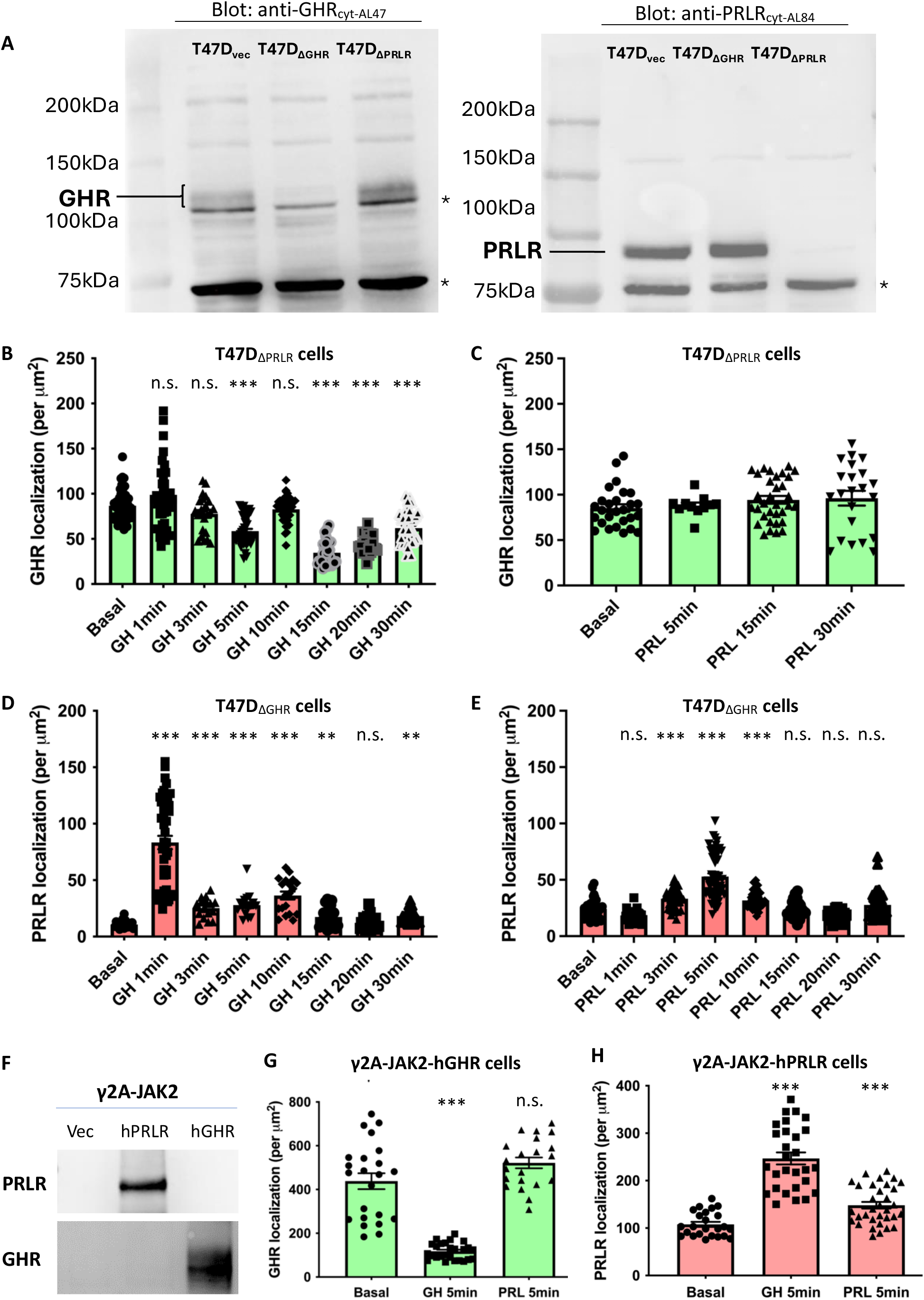
PRL-induced reduction of hGHR on the cell surface depends on the presence of PRLR. **(A)** Detergent cell extracts of T47D_vec_, T47D_ΔGHR_, and T47D_ΔPRLR_ were analyzed by immunoblotting with anti-GHR_cyt-AL47_ and anti-PRLR_cyt-AL84_ antibodies. The bands corresponding to GHR and PRLR are annotated. Asterisks indicate migration of strong non-specific bands detected in the Western Blots, which serves as a loading control for protein. **(B)** In T47D_ΔPRLR_, GH (500 ng/ml) induces downregulation of hGHR localizations on the cell surface, while **(C)** PRL does not change hGHR localizations. **(D, E)** In T47D_ΔGHR_ cells, GH (D) and PRL (E) incite an increase of hPRLR on the cell surface. Data are displayed as mean ± SE. * (P<0.05), ** (P<0.01), and *** (P<0.001) indicate the statistical significance in comparison with Basal and are calculated by a two-tailed t-test assuming unequal variance. **(F)** γ2A-JAK2 cells with stable expression of hGHR or hPRLR. Detergent extracts were resolved by SDS-PAGE and immunoblotted with anti-GHR_cyt-AL47_ and anti-PRLR_cyt-AL84_ antibodies. Vec corresponds to stable transfection with empty vector. **(G, H)** Serum-starved γ2A-JAK2- hGHR cells (G) or γ2A-JAK2-hPRLR cells (H) were treated with GH (500 ng/ml) or PRL (500 ng/ml) for 5 min. Localizations of hGHR in γ2A-JAK2-hGHR cells were decreased after GH treatment while remaining the same after PRL treatment compared to the basal state. Localizations of hPRLR in γ2A-JAK2-hPRLR cells were increased by GH or PRL treatment. Data are displayed as mean ± SE. * (P<0.05), ** (P<0.01), and *** (P<0.001) denote the statistical significance in comparison with Basal and are calculated by a two-tailed t-test assuming unequal variance.

To extend our observations, we next examined the GH and PRL responses in a cellular reconstitution system: γ2A-JAK2 [78–80] is a human JAK2-deficient fibrosarcoma cell line reconstituted with JAK2 that stably expresses JAK2 but lacks hGHR and hPRLR. To independently study the role of each receptor in this setting, we used our previously generated stable transfectants of γ2A-JAK2 cells that harbor either hGHR or hPRLR [81] and verified the presence of the indicated receptor by immunoblotting (Fig. 5F). Consistent with the observation in T47D_ΔPRLR_ cells, γ2A-JAK2-hGHR cells responded with a loss of surface hGHR density to GH stimulation but not to PRL stimulation (Fig. 5G). Interestingly, as in T47D_ΔGHR_ cells, treatment of γ2A- JAK2-hPRLR cells with either GH or PRL (Fig. 5H) promotes increased surface hPRLR. Thus, our findings in both cell systems suggest that hPRLR-hGHR interaction, direct or indirect, is indispensable for PRL-induced but not for GH-induced loss of surface hGHR.

### Box 1 region in hPRLR contributes to PRL-induced cell surface GHR downregulation

The hPRLR-dependent PRL-induced downregulation of cell surface hGHR indicates that hPRLR can modulate the density of hGHR on the cell surface in response to PRL. To investigate this interaction, we generated a set of truncation or deletion mutants of hPRLR: (1) hPRLR-tr292, which truncates the intracellular domain of hPRLR distal to the membrane-proximal intracellular domain box 1 element; (2) hPRLR-tr238, which contains only 4 amino acids (aa) of the proximal intracellular domain and does not include box 1; and (3) hPRLR-Δbox1, in which the box 1 region (243aa-251aa) is internally deleted (shown in Fig. 6A). Expression of hPRLR-tr292, hPRLR-tr238, and hPRLR-Δbox1, as well as wild-type hPRLR, was detected by immunoblotting using monoclonal antibodies targeting the ECD of PRLR (mAb_ext-1.48_). Since hPRLR-tr238 contains only 4 aa in its ICD, no immunoblot signal was detected using a polyclonal antibody (Ab_AL-84_) targeting the hPRLR ICD. In contrast, the expression of hPRLR-tr292 was easily detected by Ab_AL-84_ (Fig. 6B). We then transiently transfected each hPRLR construct into the γ2A-JAK2-hGHR cells, which stably express hGHR, and analyzed the changes of cell surface hGHR localizations. In cells expressing wild-type hPRLR (hPRLR-WT), the localizations of hGHR significantly decreased 3 min post-exposure to GH and PRL (Fig. 6C). Similarly, under the same treatment, in cells expressing hPRLR-tr292, the localization of hGHR on the cell membrane was reduced in response to each ligand (Fig. 6D). However, in hPRLR-tr238- expressing cells, the localization of hGHR was diminished by GH stimulation but slightly increased by PRL stimulation (Fig. 6E). To further investigate the role of the box1 region for hGHR and hPRLR functional interaction, we studied hPRLR-Δbox1 expressing cells and found that upon PRL treatment, hGHR localizations did not decrease on the cell surface (Fig. 6F). As a negative control, we transfected the cells with vector (pcDNA3.1). hGHR localization decreased upon GH stimulation but remained at basal levels upon PRL stimulation (Fig. 6G). In addition, in hPRLR-ΔBox1 and hGHR expressing cells, the JAK2 and STAT5 tyrosine phosphorylation levels were assessed in response to GH and PRL stimulations. Treatment of 500ng/ml GH induces a dramatic increase in both JAK2 and STAT5 phosphorylation. In contrast, 500ng/ml PRL treatment does not cause JAK2 or STAT5 phosphorylation (Fig. 6H), consistent with PRLR Box1 being required for effective coupling of PRL occupancy of PRLR to activation of JAK2 and phosphorylation of STAT5 and the inability of PRL to signal via GHR.

**Fig. 6.**
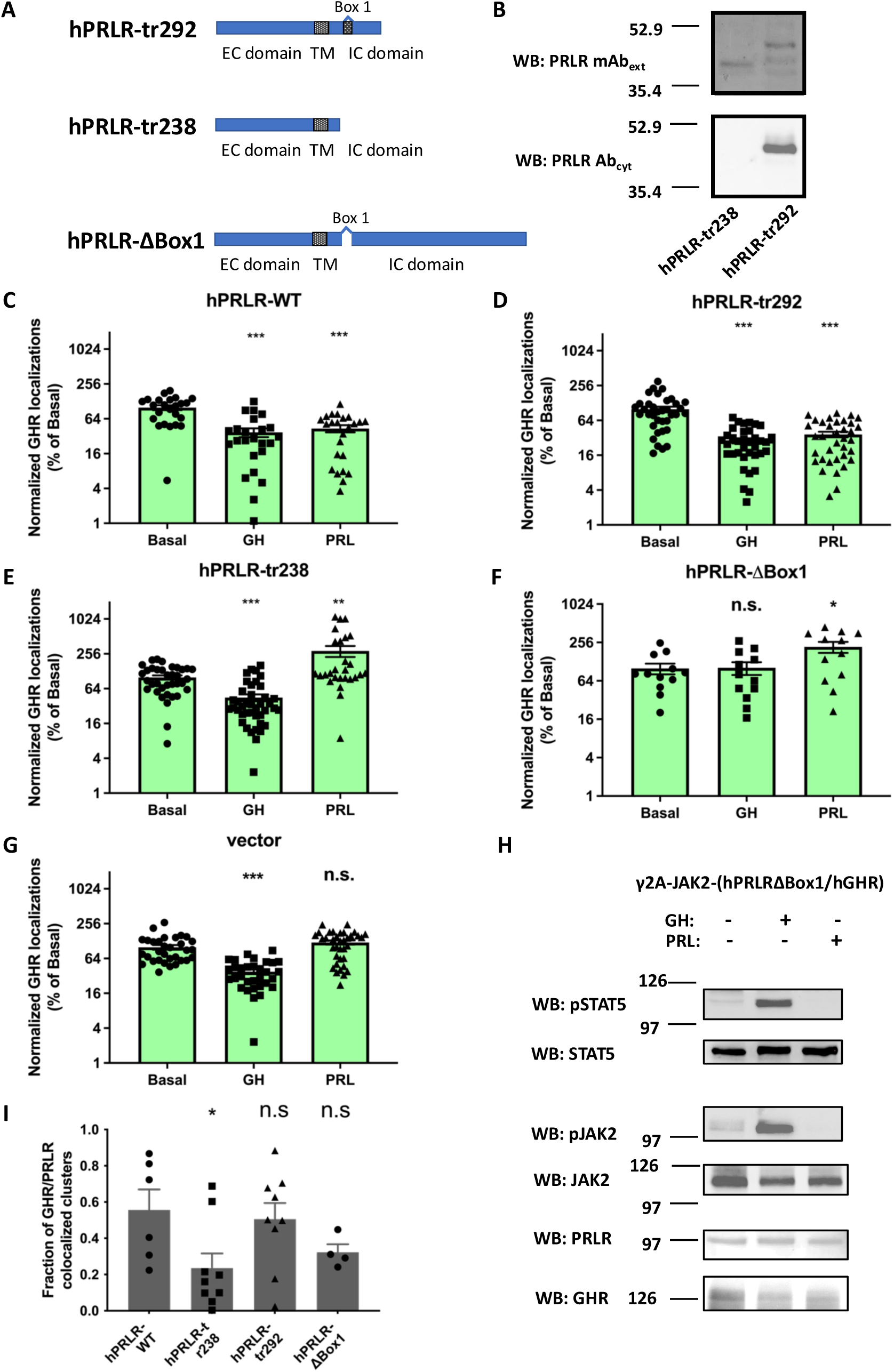
Box 1 region in hPRLR contributes to PRLR-induced downregulation of cell surface GHR. **(A)** Diagrams of employed mutant hPRLR variants with truncations or deletion in the intracellular domain. ECD, extracellular domain; TMD, transmembrane domain; ICD, intracellular domain. **(B)** γ2A-JAK2-hGHR cells were transiently transfected with hPRLR-tr292 and hPRLR-tr238. Cell extracts were resolved by SDS-PAGE and blotted with anti-PRLR mAbext-1.48 and anti-PRLRcyt-AL84. **(C-G)** γ2A-JAK2-hGHR cells were transfected with (C) wild-type hPRLR (hPRLR-WT), (D) hPRLR truncated at 292 aa (hPRLR-tr292), (E) hPRLR truncated at 238 aa (hPRLR-tr238), (F) hPRLR with box1 motif deleted (hPRLR-ΔBox1) and (G) vector pcDNA3.1 (vector). Transfected cells were imaged by dSTORM microscopy and analyzed using the DBSCAN algorithm. The density of hGHR localizations on the cell surface was calculated. Each data point represents the density of hGHR in a cell surface area of size 6.25 μm X 6.25 μm. Data are collected from at least 6 cells (4 ROIs per cell) from each group, and values are displayed as mean ± SE (from three independent experiments). Data are normalized such that the basal is 100%. * (P<0.05), ** (P<0.01), and *** (P<0.001) denote the statistical significance in comparison with Basal and are calculated by a two-tailed t-test assuming unequal variance. **(H)** Detergent cell extracts of hPRLR-ΔBox1 and hGHR-expressing cells were analyzed by immunoblotting. After 5 hrs. starvation, cells were treated with GH (500 ng/ml) or PRL (500 ng/ml) for 10 min. In each experiment, the average Basal value is considered 100%. **(I)** Fraction of GHR/PRLR colocalized clusters. In the resting state, the colocalization ratio is significantly lower for cells expressing hPRLR-tr238 than for those expressing hPRLR-WT. Each data point represents the ratio of the number of clusters, which contain both hGHR and hPRLR, to the total number of clusters on the cell surface. Data are displayed as mean ± SE. * (P<0.05) indicates statistical significance in comparison with WT and is calculated by a two-tailed t-test assuming unequal variance (all bars without an asterisk are not significant).

Next, we analyzed colocalization in each group. In the resting state, the ratios of GHR-PRLR-colocalization clusters are relatively higher in hPRLR-WT- and hPRLR-tr292-expressing cells in comparison with hPRLR- tr238- and hPRLR-ΔBox1-expressing cells (Fig. 6I). This suggests that the box 1 region in hPRLR plays a critical role in stabilizing the hGHR-hPRLR complexes in the basal state.

### Box 1 region in hGHR plays an essential role in regulating PRLR and GHR interaction

From our observations (Fig. 6), we concluded that the JAK2 binding site, i.e., the box 1 region, in hPRLR is required for PRL-induced hGHR down-regulation from the cell surface. To further assess whether the JAK2 binding site on hGHR is also essential, we generated hGHR-ΔBox1, in which the box 1 region (297aa – 305aa) was deleted (Fig. 7A). In cells expressing both hGHR-ΔBox1 and hPRLR we observed that GH does not alter the hGHR-ΔBox1 localizations on the cell surface, while PRL slightly increases hGHR-ΔBox1 localizations (Fig. 7B). Together, these results re-affirm that binding of JAK2 to hGHR is also required for hPRLR-mediated regulation of hGHR availability on the cell surface.

**Fig. 7.**
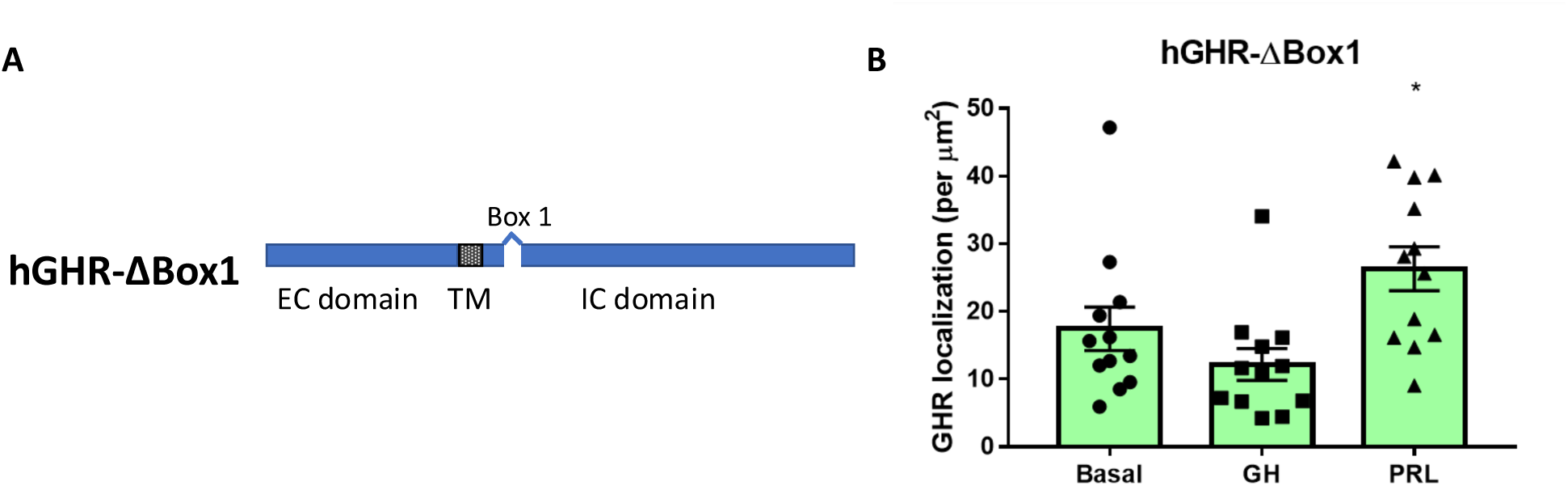
Box 1 region in hGHR also plays an important role in PRLR and GHR interaction. **(A)** Diagram of hGHR-ΔBox1 showing the deleted box 1 region in the intracellular domain of hGHR. **(B)** γ2A-JAK2-hPRLR cells were transfected with hGHR-ΔBox1. Transfected cells were imaged by dSTORM microscopy and analyzed using DBSCAN. The density of hGHR localizations on the cell surface was calculated. Each data point represents the density of hGHR in a cell surface area of size 6.25 μm X 6.25 μm. Data are collected from at least 6 cells for each group and displayed as mean ± SE. * (P<0.05), ** (P<0.01), and ***(P<0.001) indicate the statistical significance in comparison with Basal and are calculated by a two-tailed t-test assuming unequal variance.

## Materials and Methods

### Materials

Common reagents were purchased from Sigma Aldrich Corp. (St. Louis, MO) unless otherwise noted. Fetal bovine serum was purchased from Atlanta Biologicals (Lawrenceville, GA). Cell culture medium, penicillin/streptomycin and trypsin were purchased from Corning (Corning, NY). Recombinant hGH was kindly provided by Eli Lilly & Co. (Indianapolis, IN). Recombinant human PRL was obtained from the National Hormone and Pituitary Program.

### Antibodies

Polyclonal anti-GHR_cyt-AL47_ (1:1000) against the intracellular domain of GHR and polyclonal anti-PRLR_cyt-AL84_ against the human PRLR intracellular domain described previously in [82] and [83], respectively, were used as the primary antibodies for western blot analysis. Monoclonal anti-GHR_ext-mAb_ 74.3 (1:1000) against the extracellular S2 domain of GHR [40, 68, 70, 79, 81, 84–93] and monoclonal anti-PRLR_ext-mAb_ 1.48 against the extracellular S2 domain of human PRLR [70] were used for microscopy. Specifically, we let hybridoma cells secrete these antibodies into the cell medium to avoid the purification step. A detailed description of the antibodies and their previous use is provided in the references. Specificity of anti-GHR_ext-mAb_ and anti-PRLR_ext-mAb_ 1.48 was confirmed using confocal microscopy (see also Supp. Fig. S2-S4). The following secondary antibodies were used: Alexa Fluor 568 goat anti-mouse IgG1 (Invitrogen #A-21124) (1:1000), Alexa Fluor 647 goat anti-mouse IgG2b (Invitrogen #A-21242) (1:1000), Alexa Fluor 647 goat anti-mouse IgG1 (Invitrogen #A-21240) (1:1000) and Alexa Fluor 568 goat anti-mouse IgG2b (Invitrogen # A-21144) (1:1000). In the absence of primary antibodies, secondary antibodies did not show any fluorescence signals (see also Supp. Fig. S3).

### Cloning and constructs

The human GHR cDNA in pcDNA1 was a generous gift from R. Ross (University of Sheffield, Sheffield, UK). The human PRLR cDNA in pEF/V5/HIS was generously provided by C. Clevenger (Virginia Commonwealth University, Richmond, VA). hPRLR-tr238 and hPRLR-tr292 were generated by amplifying with external primers containing EcoRI site and a stop codon with XhoI site after the sequence of 238 or 292 amino acid, respectively. hPRLR-ΔBox1, and hGHR-Δbox1 were generated by overlap extension polymerase chain reaction (PCR) with associated primers and cloned into pcDNA3.1(+) vector.

### Cell culture and transfections

Human T47D breast cancer cells were purchased from American Type Culture Collection (Manassas, VA). Cells were maintained in RPMI 1640 medium supplemented with 10% fetal bovine serum (FBS), 100 units/ml penicillin and 100 μg/ml streptomycin in a humidified atmosphere of 5% CO2 and 95% air at 37 ℃.

γ2A-JAK2 cells were generated by transfection of γ2A cells [78] with pcDNA3.1(+)/zeo-JAK2 and maintained in culture, as described previously [79, 80]. The generation of γ2A-JAK2-GHR and γ2A-JAK2- PRLR cells has been previously described [81].Transient expression of receptors was achieved by using Lipofectamine LTX Plus (Invitrogen), transfecting 0.3 pmol plasmid DNA per 6-cm^2^ dish.

### Western blot

Cells were serum starved for 5 hours and treated with 500 ng/ml GH or 500 ng/ml PRL at 37 °C for 10 min. Stimulations were terminated by washing the cells with ice-cold phosphate-buffered saline supplemented with 0.4 mM sodium orthovanadate. Cells were lysed in lysis buffer for 30 min at 4°C. Then cell lysates were centrifuged at 15,000 g for 10 min at 4°C. The protein extracts (supernatant) along with SDS sample buffer were resolved by SDS-PAGE.

### Sample preparation for imaging

Cells were seeded into either eight well #1.5 coverslip bottom dishes (Ibidi) or 25 mm #1.5 coverslips (Electron Microscopy Sciences). Cells were serum starved for 5 hours, then treated with GH (500 ng/ml), PRL (500 ng/ml) at 37 °C for the indicated time. Then the cells were washed with Dulbecco’s Phosphate-Buffered Saline (DPBS) three times and fixed with 4% PFA (paraformaldehyde) in 1X DPBS with calcium and magnesium (Cat # 21-030-CV) for 10 min. After multiple washes with DPBS, cells were blocked with 5% normal horse serum, 5% normal goat serum, 5% BSA for 30 min. Lastly, cells were incubated in primary antibody for 3h and secondary antibody for 70 min at 37°C.

### dSTORM imaging and data analysis

dSTORM experiments were conducted on an inverted Nikon Ti2 N-STORM microscope equipped with a 100 × 1.49 NA oil immersion objective (Nikon, Japan), 488 and 647 nm lasers, and an iXon DU-897 Ultra EMCCD camera (Andor, Oxford Instruments). The dSTORM imaging buffer consists of two buffers, A and B, and a GLOX mixture. Buffer A is made of 10 mM Tris pH 8 and 50 mM NaCl, while buffer B is made of

50 mM Tris pH 8 and 10 mM NaCl. The GLOX mixture consists of 14 mg glucose oxidase (Sigma, St Louis, Missouri), 50 μl catalase (17 mg/ml) (Roche, Penzberg, Germany), and 200 μl of Buffer A. The dSTORM buffer is made by combining (on ice) 7 μl GLOX with 7 μl 2-mercaptoethanol (Sigma, St Louis, Missouri) and 690 μl Buffer B, using only the supernatant from the GLOX mixture. For a detailed description of the buffer, we refer the reader to [94].

The cell membrane was focused using TIRF excitation, which selectively images within 100-150 nm of the cell membrane, making it an excellent method to study surface distribution of membrane proteins. Of note, TIRFM exploits the unique properties of an induced evanescent field in a limited specimen region immediately adjacent to the interface between two media having different refractive indices. It thereby dramatically reduces background by rejecting fluorescence from out-of-focus areas in the detection path and illuminating only the area right near the surface.

Moreover, we reduced auto-fluorescence by ensuring that (a) cells are grown in phenol-red-free media, (b) imaging is performed in STORM buffer which reduces autofluorescence, and (c) our immunostaining protocol includes a quenching step aside from using blocking buffer with different serum, in addition to BSA (5%). Moreover, we employed extensive washing steps following antibody incubations to eliminate non-specifically bound antibodies. Ensuring that the TIRF illumination field is uniform helps reduce scatter. Additionally, an extended bleach step prior to the acquisition of frames to determine localizations helped further reduce the probability of non-bleached fluorescent molecules.

For dual color STORM imaging, the sample was illuminated with alternating 561 nm or 647 nm lasers and 40,000 frames of images were acquired per channel (9ms/frame). The chromatic offset associated with the different acquisition wavelengths was corrected using 0.1 μm microspheres [95, 96]. Imaging captured the fluorophores’ blinking events. Localizations were counted as true localizations, when at least 5 consecutive blinking events had been observed. Moreover, coordinates were filtered using a threshold radius of 25nm. dSTORM images were reconstructed using the built-in STORM module in NIS-Elements (Nikon). Nikon software was used for Gaussian fitting, and each dot represents the peak of a Gaussian. The localizations list was exported and further analyzed in *Matlab*. In particular, for the DBSCAN cluster analysis, we employed Clus-DoC directly on the list of localizations, utilizing the parameters of a minimum of 4 neighbors within a cluster radius of 20nm [97].

### Proximity Ligation Assay

Cells were serum starved for 5 hours, then treated with GH (500 ng/ml) or PRL (500 ng/ml) at 37 °C for the indicated time. Then the cells were washed with DPBS three times and fixed with 4% PFA (paraformaldehyde) for 10 min. After multiple washes with DPBS, cells were blocked with 5% normal horse serum, 5% normal goat serum, 1% BSA for 30 min. Lastly, cells were incubated in primary antibody for 3h and secondary antibody for 70 min at 37°C. After secondary antibody incubation, ligation and amplification were performed according to the manufacturer’s protocol (Duolink® PLA; Sigma-Aldrich). Briefly, ligation solution containing ligase was added to the samples, followed by incubation for 30 minutes at 37°C. Samples were then washed and incubated with the amplification solution, which includes polymerase, at 37°C for 100 minutes to amplify the PLA signal.

### Generating hGHR knockout and hPRLR knockout T47D cells

We utilized CRISPR/Cas9 to generate T47D_ΔPRLR_ and T47D_ΔGHR_. Two single-guide RNAs (sgRNAs) and Cas9 nickase (Cas9n) were designed to create sticky-end double-strand breaks in DNA at the targeted locus, triggering DNA repair via the Non-Homologous End Joining (NHEL) pathway, which introduces insertions, deletions or frameshift mutations [98]. Each sgRNA was expressed using plasmid #62987 pSpCas9n(BB)- 2A-Puro(PX462) V2.0 (purchased from Addgene), which also expressed Cas9n when transfected into cells. Two vectors, built on pSpCas9n(BB)-2A-Puro(PX462) V2.0 backbone, containing a pair of sgRNAs targeting either the hGHR or hPRLR gene for knockout, as well as a built-in Cas9n gene, were transfected into newly purchased T47D cells. The transfected cells were plated at serial dilutions and selected with puromycin. Single clones were picked and expanded. The vector backbone pSpCas9n(BB)-2A-Puro(PX462) V2.0 was also used for transfection into T47D cells as a control cell line.

Genomic DNA was extracted from each clone. Primers were designed 200-400 bp upstream and downstream of the targeted locus of the hGHR or hPRLR gene to amplify the genomic DNA via PCR, which was subsequently cloned into the pUC19 vector for sequencing. Clones with mutations on both alleles were identified, and the western blot was used to confirm the absence of expression of the targeted genes.

### Design of sgRNAs for hGHR or hPRLR knockout

A pair of 20 bp sequences located 10-25 bp apart at the target locus was chosen using the Feng Zhang lab’s Target Finder [99]. This tool identified the best pair of guide RNA sequences with theoretically zero off-target effects. One sequence targeted the top strand, and the other targeted the bottom strand, enabling the introduction of a sticky-end double-strand DNA break. Each guide RNA sequence was synthesized as double-stranded oligonucleotides with a CACC at 5’ overhand of the 20-nt guide sequence and an AAAC at the 5’ overhand of the complementary strand, allowing insertion into the backbone plasmid pSpCas9n(BB)-2A-Puro(PX462) V2.0 using the Bbsl restriction enzyme. An extra nucleotide ‘G’ was added to the 5’ end of the 20-nt guide if the sequence did not begin with ‘G’, which is the preferred first base following the U6 RNA Polymerase III promoter.

The target locus for introducing mutations in the hGHR gene was located at exon 4. Exon 4 was chosen because many naturally occurring pathological mutations in Laron Syndrome are located in this region. Additionally, deletion of the 3’ end of exon 4 and its adjacent intron has been shown to successfully generate GHR knockout mice[100]. The hPRLR gene has seven isoforms; exon 5 is included in six of these isoforms, except for the ΔS1 PRLR isoform, in which both exons 4 and 5 are skipped by alternative splicing. Therefore, we selected the end of exon 5 and the adjacent intron as the target locus for introducing mutations. The goal was to knock out all six isoforms containing exon 5, as well as the ΔS1 PRLR isoform, by interrupting alternative splicing. This was achieved by deleting the initial GU dinucleotide sequence at the 5’ end of the intron, which is required for the binding of the U1 protein of the spliceosome.

## Statistical analysis

Imaging and biochemical experiments were carried out at least three times to ensure reproducibility. Data were analyzed using unpaired, two-tailed t-tests. *Prism* software was used for statistical analysis (GraphPad Inc, USA). Data are presented as means ± s.e.m.

To quantify changes in localization and colocalization we randomly selected 4 ROIs (regions of interest) per cell to calculate fractions and then calculated the average of three different cells from independently repeated experiments.

## Discussion

The non-ligand-bound states of hGHR and hPRLR have been extensively investigated. Pre-homodimerization of hGHR and hPRLR has been reported using structural and biochemical methods [40, 101–104]. At the same time, a recent study supports the notion that GHR, at physiological densities, exists as monomers on the cell surface and becomes activated by a ligand-induced dimerization [105]. The current study uses a highly precise and sensitive single-molecule localization microscopy approach to study the formation of hGHR and hPRLR nanoclusters and the cell surface receptor availability. Our super-resolution images reveal that in human T47D breast cancer cells and the γ2A-JAK2 cell exogenous expression system, the cluster size of hGHR and hPRLR in the basal state range from a few localizations per cluster up to a thousand localizations per cluster.

With GH or PRL treatment, the number of hGHR on both T47D and γ2A-JAK2 cell surfaces is decreased, indicating the removal of surface hGHR. Moreover, the distribution curve of hGHR shifts toward larger cluster sizes (Fig. 3), suggesting a ligand-induced aggregation of receptors. In turn, hPRLR numbers dramatically increase on the cell surface in response to ligand stimulation. The newly presented hPRLR clusters may impact the distribution of cluster sizes, which may explain why the median of hPRLR cluster sizes does not change much with ligand treatment.

Our PL assay data indicate that distances between hGHR and hPRLR are small (less than 40nm). This implies that hGHR and hPRLR form co-localized clusters in unstimulated states, suggesting hGHR and hPRLR are physically accessible to one other. With GH stimulation, the fraction of co-localized receptors is decreased. Given that GH treatment enhances the coimmunoprecipitation of total cellular hGHR and hPRLR [68], the decrease in co-localized surface receptors can likely be attributed to removing co-localized receptors from the cell surface.

It is well documented that hPRLR is engaged by both PRL and GH, while hGHR only responds to GH [41, 43, 44]. Unexpectedly, we found that PRL induces a down-regulation of cell surface hGHR in cells that co-express hGHR and hPRLR (Fig. 2), indicating that hPRLR directly or indirectly interacts with hGHR. Interestingly, PRL was unable to induce hGHR down-regulation in cells either lacking hPRLR or expressing hPRLR without its Box1 region (hPRLR-tr238 or hPRLR-ΔBox1), suggesting that JAK2 and PRLR association is required for hGHR-hPRLR interaction that in turn allows PRL-induced downregulation of cell surface hGHR. Notably, previous work showed that GHR and JAK2 association is necessary for JAK2 to stabilize cell surface GHR and inhibit constitutive GHR down-regulation [80]. Furthermore, single-particle tracking studies showed that, in the presence of JAK2, a higher level of ligand-induced dimerization of GHR was observed [105]. Moreover, JAK2 with intact kinase activity is required for GH-induced GHR down-regulation [80]. Similarly, the box1 region of PRLR also associates with JAK2, and deletion or modulation of the last proline residue of box1 abrogates PRLR function [106]. Collectively, these findings suggest that the box1 regions in hGHR and hPRLR play a crucial role in the transduction of the individual signaling cascades and hGHR-hPRLR association.

Here we found that, in the resting state, the degree of hGHR-hPRLR colocalization is higher in cells expressing hPRLR with the box1 motif. This suggests JAK2 may not only stabilize GHR but also support the formation of hGHR-hPRLR-containing clusters. Previously, it has been suggested that such clusters are comprised of hGHR homodimers and hPRLR homodimers that together form (hGHR-hGHR) – (hPRLR- hPRLR) hetero-multimers or higher order oligomers [69]. Indeed, because the intracellular domains (ICD) of both hGHR and hPRLR are highly disordered, the flexibility of their ICDs may provide room for recruiting JAK2 and stabilizing the hGHR-hPRLR association [107, 108]. Moreover, a study of the crystal structure of erythropoietin receptor (EPOR) and leptin receptor (LEPR), which also belong to the class I cytokine receptor family, revealed recently that JAK2/EPOR and JAK2/LEPR complexes contained four JAK2 and four EPOR or LEPR molecules, respectively [109]. Hence, it is possible that hGHR-hGHR homodimers and hPRLR-hPRLR homodimers form complexes in a similar fashion (Fig. 8). We propose that this JAK2/Box1-mediated interaction of receptors is not limited to hGHR and hPRLR but may generalize to other receptors of the class I cytokine receptor family. We note, however, that determinants within particular receptors (perhaps residing in their extracellular and/or transmembrane domains) and their intracellular JAK2 association domains may facilitate, to varying degrees, their propensity to form multimeric aggregates. GHR and PRLR, for example, may tend to do so more avidly with each other than either one does with other cytokine receptors. Additional studies are required to determine and understand the modes and functions of the hGHR-hPRLR association.

**Fig. 8.**
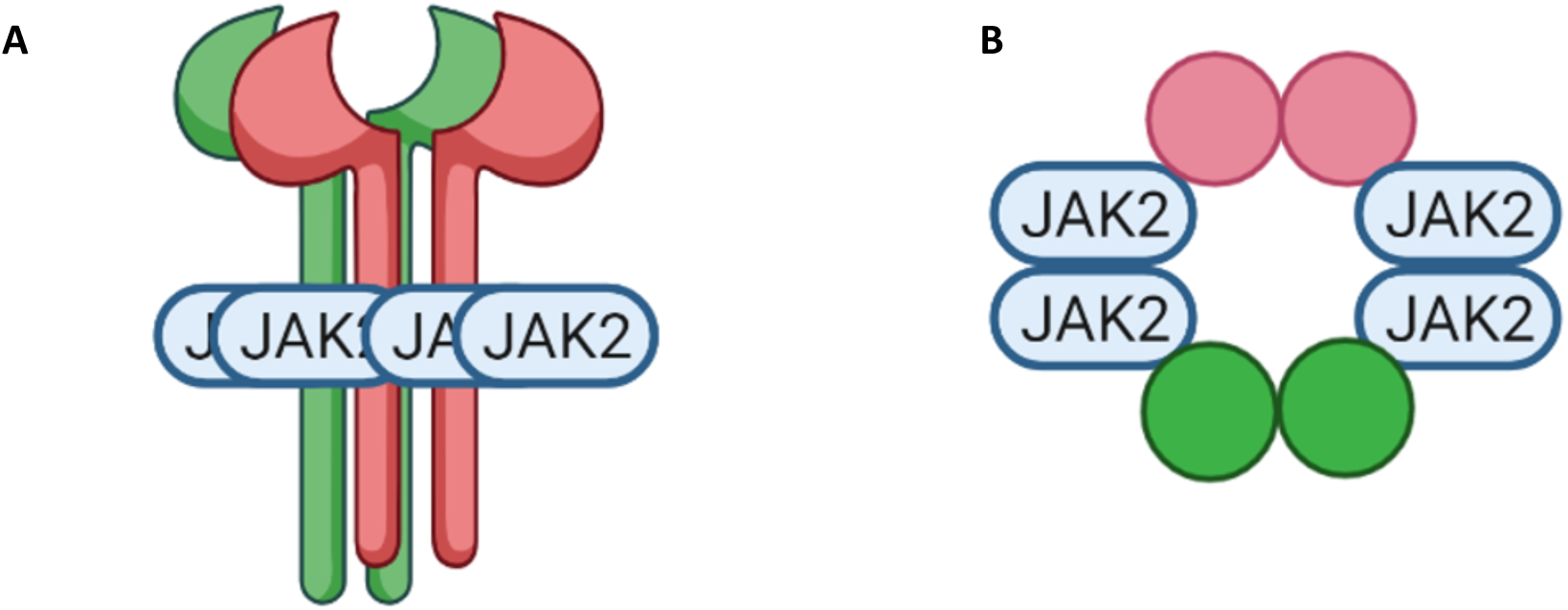
Model of JAK2 stabilizing GHR-GHR and PRLR-PRLR heteromers. **(A)** Schematic representation of GHRs homodimer and PRLRs homodimer form a heteromer, which contains two PRLRs (red), two GHRs(green), and four JAK2(blue). **(B)** A top view of the JAK2/PRLR/GHR multimer.

An obvious question is to what extent our observed regulation is biologically relevant. While our work does not provide an answer to this question, one could argue that, generally speaking, such a mechanism could provide a means for crosstalk and feedback between the two cytokine receptors and allow for a more fine-tuned signaling response. In particular high levels of PRL could reduce GH signaling thereby prioritizing PRL’s function over GHR-mediated growth processes. PRLR’s ability to modulate GHR cell surface availability might also contribute to tissue-specific responses to hormonal signals. For instance, in situations with high PRL levels in circulation such as during lactation [110], downregulation of mammary gland cell surface GHRs could prevent GH from counteracting PRL’s effects. Also, PRL and GH have important roles in metabolism, often with opposing effects [2, 111–113]. PRL promotes lipogenesis and energy storage, while GH promotes lipolysis and energy mobilization. By downregulating GHRs, PRL could shift the metabolic balance in favor of energy storage. Moreover, this regulation may also have disease implications: hPRL/hPRLR-mediated downregulation of cell surface hGHR could be a relevant factor in the observed anti-tumor effect of PRLR signaling, especially in triple-negative breast cancer (TNBC) expressing PRLR but lacking other interfering hormone receptors. As previously shown, the TNBC cell lines MDA-MB- 231 and MDA-MB-453 show significantly increased GHR [15] while expressing no or only endogenous levels of PRLR, respectively [28]. PRL treatment significantly decreased cell viability and invasive capacity of MDA-MB-231 cells following the restoration of PRLR expression. In MDA-MB-453 cells, PRL caused a significant reduction in cell viability. A xenograft model with inoculated MDA-MB-453 cells confirmed the growth inhibitory effect of PRL treatment *in vivo*, suggesting PRLR expression as an indicator of a favorable prognosis. However, several studies have reported the contrary, namely that PRL promotes breast cancer [12], presenting a conundrum that so far has not been resolved. Among the observations supporting a role of PRL in promoting breast cancer are: (1) frequent PRLR overexpression in BC tissues, (2) association of elevated (serum) PRL with increased BC risk – in particular for estrogen receptor (ER) positive BC – and increased risk for metastatic progression from epidemiologic data, and (3) PRL stimulation modulating the cytoskeleton and inducing motility of human breast cancer cells (including T47D) in vitro, which has been associated with mammary carcinoma progression in vivo [12]. Of note, our hypothesis of PRL-mediated down-regulation of GHR presence on the cell surface being protective does not mean that other effects cannot also be involved or cancel this positive effect out. Moreover, timing could be a decisive element: while PRL might have a protective effect (depending on other factors) at the time of tumorigenesis, it may actually facilitate BC metastasis. But given the lack of experimental data indicating a clear protective effect through Jak2-mediated receptor interactions in PRLR+ GHR+ BC at any BC stage, our hypothesis remains speculative.

Besides, observations in MCF-7, a human breast cancer cell line, where binding of PRL causes internalization and downregulation of PRLR expression seem to contradict our observations in T47D cells [114]. Recently, a competition between two kinases (JAK2 and LYN) for binding hGHR at the box1-box2 region was reported [115], indicating that hGHR nanocluster formation at the cell membrane is regulated also by kinases. It is conceivable that similar kinase-dependent competition mechanisms could regulate also PRLR cell surface availability. Thus, differences in cells’ expression of such kinases could then play a role in the perceived inconsistency. Moreover, [114] focuses on the downregulation of the long PRLR isoform in response to PRL, while all other isoforms were undetectable in MCF-7 cells. So, this and other differences between the cell lines may be the reason for the perceived contradiction.

We note that a limitation of our study is the focus on T47D and γ2A-JAK2 cells. It remains to be seen, if we can observe and replicate the same effect also in other cell lines, for instance mammary epithelial cells or other hGHR+ PRLR+ cancer cell lines. Also, the roles of the different membrane PRLR isoforms have not been fully addressed and our study lacks isoform specific data [37]. The anti-PRLR antibody that we used does not only recognize the long form PRLR but also the intermediate form and the short forms, all of which have ECDs. All these forms contain the box1 motif. Importantly, form S1b is almost identical to hPRLR-tr292, but behaves similarly to LF (i.e. our hPRLR WT) in terms of normalized GHR cell surface localization and PRLR co-localization (Fig. 6). So, while we do not have experimental confirmation, we would expect that these other isoforms behave somewhat similarly to LF with respect to GHR surface membrane interactions – while still inducing different downstream effects as well as playing different roles in breast cancer [37, 116–118].

As part of future research, we plan to study the observed effect in cell lines other than T47D. We would also like to investigate if the mechanism is PRLR-GHR specific, or if there are other cytokine receptors that can interact in a similar asymmetric, ligand-dependent manner, such that one regulates the other receptor’s surface availability. Importantly, we would like to elucidate any biological relevance of the observed effect, analyze roles and consequences of PRLR isoforms, and investigate our hypothesis that PRLR-mediated downregulation of cell surface GHR contributed to the anti-tumoral effect of PRL treatment in PRLR-positive TNBC.

## Notes

### Competing Interest Statement

The authors have declared no competing interest.

### Summary of Updates

We have significantly revised the manuscript. The introduction now includes also information on different PRLR isoforms as well as alternative JAK2 signaling pathways. The Methods section includes now new subsections on PLA, the generation of hGHR/hPRLR knockout T47D cells, and the design of sgRNAs. Information on used antibodies was updated. We have also added more information on dSTORM imaging and the dSTORM buffer. The discussion includes four new or extended paragraphs discussing biological relevance, perceived contradictions to previous reports, limitations of our study, and future research. Figures and/or figure legends were updated. Fig. 1 now includes zoomed-in sections. Fig 5 shows larger crops of WBs. Data of controls indicating antibody specificity (using confocal microscopy) have been added to the manuscript's supplementary material (see Figs. S2 to S4).

