## Supplementary Material for "A role for JAK2 in mediating cell surface GHR-PRLR interaction"

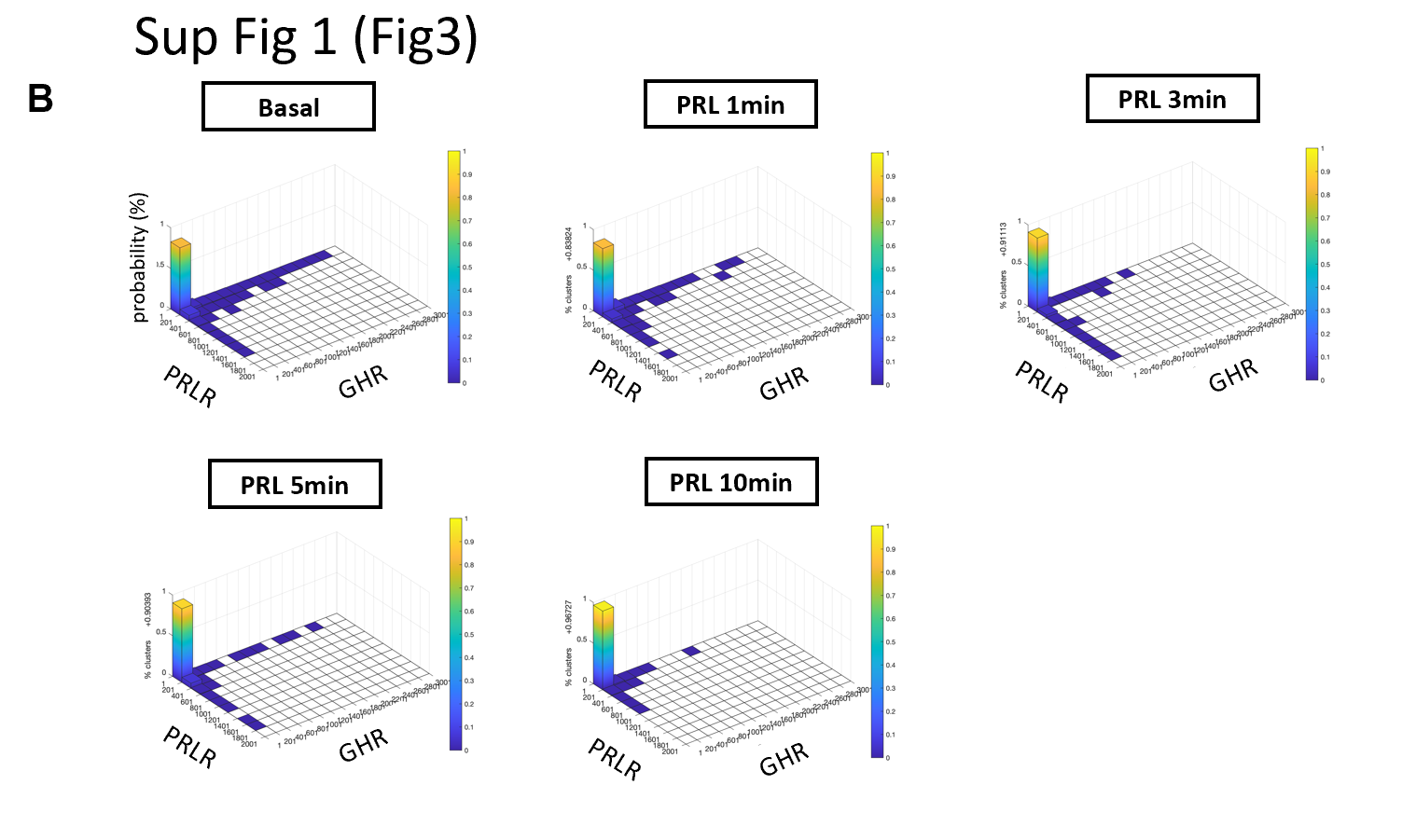
Supplementary Figure:

**B**

**A**

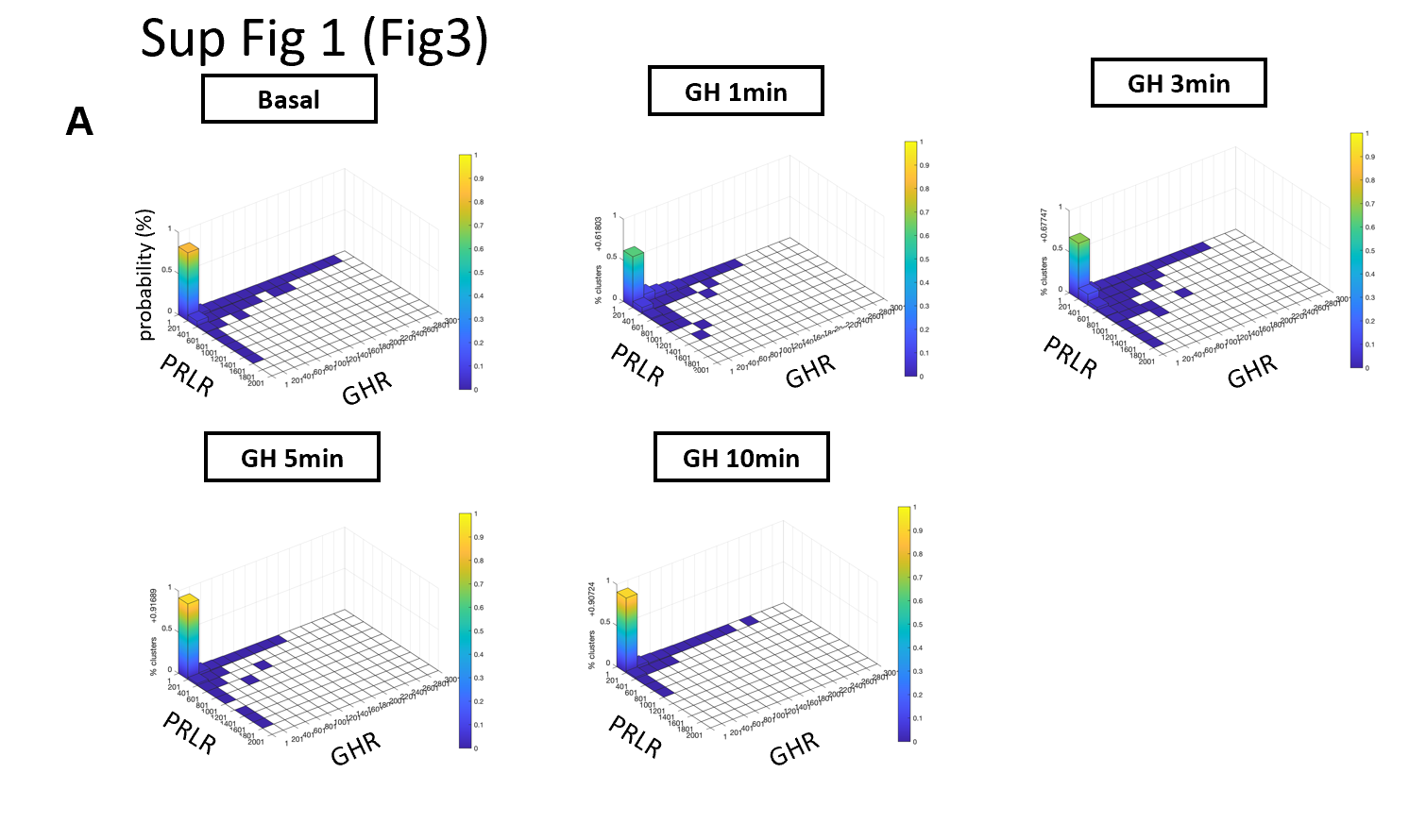

**Fig. S1:** The size distributions of co-localized clusters are shown in 3D plots, where the binned x-axis represents the number of hGHR localizations, the binned y-axis represents the number of hPRLR localizations, and the z-axis represents the probability of the cluster. Each bin has a size of 200 localizations. Both GH **(A)** and PRL **(B)** induce a distinct change in the composition of co-localized clusters.

**Fig. S2:** Confocal microscopy images indicating primary antibody specificity. Rows 1 & 3: Images of γ2A-JAK2 cells stably expressing GHR (no PRLR). Rows 2 & 4: Images of γ2A-JAK2 cells stably expressing PRLR (no GHR). Rows 1 & 2: Cells were incubated in hybridoma medium secreting primary GHR 74.3 (IgG1) antibodies. Rows 3 & 4: Cells were incubated in hybridoma medium secreting primary PRLR 1.48 (IgG2) antibodies. As secondary antibody we used antimouse-Alexa 568. In all images, brightness was increased by 40%.

**Fig. S3:** Confocal microscopy images indicating secondary IgG2b antibody specificity. Secondary antimouse (IgG2b) antibodies conjugated with Alexa 568 (1^st^ row) and Alexa 647 (2^nd^ row) fluorophores bind primary antibody PRLR-Ab_ext_ 1.48 (right two columns) but not GHR-Ab_ext_ 74.3 (left two columns). In all images, brightness was increased by 40%.

**Fig. S4:** Confocal microscopy images demonstrating absence of unspecific secondary antimouse-Alexa 568 binding. In all images, brightness was increased by 40%.

Supplementary Methods:

1. **The synthesized oligonucleotides for guide RNAs.**

GHR guide A

GHR guide A top	5’- CACC**G** TCGCTCAGGTGAACGGCACT – 3’

GHR guide A bottom	3’ - **C** AGCGAGTCCACTTGCCGTGA CAAA – 5’

GHR guide B

GHR guide B top	5’- CACC**G** TGGACAGATGAGGTTCATCA – 3’

GHR guide B bottom	3’ - **C** ACCTGT CTA CTCCAAGTAGT CAAA – 5’

PRLR guide A

PRLR guide A top	5’- CACC**G** CCGAGAAACTGCTTCCCATC – 3’

PRLR guide A bottom	3’ - **C** GGCTCTTTGACGAAGGGTAG CAAA – 5’

PRLR guide B

PRLR guide B top	5’- CACC**G** GACTTACATAGGTAAGGGGA – 3’

PRLR guide B bottom	3’ - **C** CTGAATGTAT CC ATTCC CCT CAAA – 5’
